# Generating novel protein sequences using Gibbs sampling of masked language models

**DOI:** 10.1101/2021.01.26.428322

**Authors:** Sean R. Johnson, Sarah Monaco, Kenneth Massie, Zaid Syed

## Abstract

Recently developed language models (LMs) based on deep neural networks have demonstrated the ability to generate fluent natural language text. LMs pre-trained on protein sequences have shown state of the art performance on a variety of downstream tasks. Protein LMs have also been used to generate novel protein sequences. In the present work we use Gibbs sampling of BERT-style LMs, pre-trained on protein sequences using the masked language modeling task, to generate novel protein sequences. We evaluate the quality of the generated sequences by comparing them to natural sequences from the same family. In particular, we focus on proteins from the chorismate mutase type II family, which has been used in previous work as an example target for protein generative models. We find that the Gibbs sampling process on BERT-style models pretrained on millions to billions of protein sequences is able to generate novel sequences that retain key features of related natural sequences. Further, we find that smaller models fine-tuned or trained from scratch on family-specific data are able to equal or surpass the generation quality of large pre-trained models by some metrics. The ability to generate novel natural-like protein sequences could contribute to the development of improved protein therapeutics and protein-catalysts for industrial chemical production.

## 1. Introduction/Background/Motivation

### 1.1. Background

Proteins are critical components of biological systems, contributing to many functions within cells, including cellular structure, regulation of gene expression, protection against pathogens, and catalysis of chemical conversions. In 2018, the Nobel Prize in Chemistry was awarded to Frances Arnold for her work in directed evolution, the iterative improvement of protein function by making small changes to a starting sequence in a step-wise process. Directed evolution experiments have produced new proteins with improved or altered functions [6, 2].

Proteins are made up of building blocks called amino acids. There are 20 different amino acids, which are commonly referred to by one-letter abbreviated names. Amino acids in a protein are linked in a linear fashion, so a protein can be represented by a string of amino acid abbreviations, for example: ”MTSENPLLALR”. In biological systems, the linear chains fold in on themselves to form threedimensional structures where amino acids that are distant from each other in the linear sequence can be adjacent in physical space and can interact with each other. Proteins range in size from fewer than 100 amino acids up to several thousand, with most proteins having a few hundred. The total number of possible protein sequences is vast. Considering just proteins of length exactly 100, there are 20^100^ possible sequences. However, protein space is also sparse in that only a small fraction of the total number of possible proteins will fold into stable, functional conformations. Recent technological advances have made it possible to create large databases of naturally occurring protein sequences, for example [UniProt](https://www.uniprot.org/), which contains 209 million sequences, and BFD [26], which contains 2.1 billion protein sequences.

Generating and screening protein variants for directed evolution workflows is an expensive and time-consuming process. Each candidate variant has to be created in the laboratory and tested for function. Depending on the properties of the protein and the available equipment, the numbers of variants that can be screened at a time may vary from 10s to 100,000s. Given the sparsity of stable variants in protein space, most random changes to a stable protein tend to result in unstable, useless proteins. Computational methods for pre-screening candidate protein sequences, or for generating protein sequences that are likely to be stable, can improve the efficiency of directed evolution workflows by allowing experimenters to focus their efforts on sequences with the highest chances of being stable and functional.

Currently dominant methods of computational prediction of protein stability include structure-based models, supervised machine learning (ML) models, and evolution-based models. Structure-based models involve simulations of the chemical bonding between amino acids in 3-dimensional space; these models are computationally intensive and require information about the folded conformation of the protein, which is not always available. For supervised ML methods, a first round of experimental screening is done on a large and diverse set of variants, the results of the first screen are used to train parameters for a classification or regression model, and the model is used to pre-screen variants before the next round of experiments; limitations of these models are that the accuracy of these models are highly dependent on the size and quality of the training set and some models can only predict the effects of combinations of variants found in the training set.

The hypothesis behind evolution-based models is that for an organism to survive, the proteins it makes have to mostly be stable. Since the proteins in sequence databases are from living organisms, most of them presumably represent stable proteins. So a protein variant with a distribution of amino acids that resembles the distribution found in natural proteins is likely to be a stable variant. A limitation of evolution-based models is that most of the algorithms can only make useful inferences from sets of natural proteins that are closely related to the protein of interest, these proteins are said to be in the same family. Some protein families do not have very many members or their members are not very diverse, which limits the possibility to make robust statistical models from those families.

### 1.2. Related work

Recently, deep learning has been used to augment, and in many cases integrate, the computational methods mentioned above. For example, trRosetta [29] integrates structure-based and evolution-based methods by processing natural sequences using a convolutional neural network (CNN) to generate constraints for a structure based model. The task of unsupervised pre-training of language models (LMs), such as recurrent neural network (RNN) based LMs or transformer-based LMs, can readily be adapted to train on a corpus of protein sequences, such as the UniProt database, instead of a corpus of human language text. In LM pre-training on protein datasets, each protein is considered as a sentence, and each amino acid as a word. Compared to LMs for human language, LMs for protein sequences have a much smaller vocabulary, just 20 amino acids plus some control tokens, but need to be able to account for much longer sentences since protein sequences can be thousands of amino acids long. LMs pre-trained on a protein corpus (protein LMs) can be used to augment supervised learning methods, either by fine-tuning the LM for a particular downstream task or by using the model to convert sequences into sequence embeddings, then using the embeddings as features for a supervised ML model [20]. Protein LMs can also be used as a kind of drop-in replacement for ”evolution-based” models, since pre-training tasks such as next-token prediction or masked language modeling (MLM) assign probabilities to each token, the probability for the entire sequence can be determined by multiplying out the probabilities for each token in the sequence, so relative probabilities can be assigned lists of candidate protein sequences.

An example of a protein LM is UniRep [1] which is an RNN model trained on a subset of UniProt; one key finding from the UniRep work is that using the pre-training task (in their case, next token prediction) to fine-tune on additional examples of proteins from a particular family, increases performance on down-stream tasks for that family [1, 5]. Rives *et al*. [24] created 6, 12, and 34 layer models, which they call Evolutionary Scale Modeling (ESM), based on the BERT transformer encoder architecture [11] pre-trained using the MLM task. They trained ESM on the entire UniProt dataset and on subsets that were thinned to more evenly cover protein space and reduce oversampling. Rives *et al*. found that increasing model size improved performance on downstream tasks, but that the thinned training data led to better performance than the full dataset. Elnaggar *et al*. [12] pre-trained a variety of transformer-based LMs on protein sequence data including BERT, XLNet [30], Albert [15], and Transformer-XL [10]. They trained their models on the entire UniProt dataset, and on an even bigger dataset, [BFD](https://bfd.mmseqs.com/); similar to ESM, they found that bigger training datasets did not improve performance by much, which might mean that alternative model architectures or pre-training tasks may be necessary to further improve model performance. Of the architectures they tested, they found that their BERT model performed better than the others, suggesting that the transformer encoder architecture trained on the MLM task is able to capture important features of proteins more effectively than other existing transformer architectures and pretraining tasks.

Protein LMs can be used directly for protein sequence generation: Madani *et al*. [17], trained a transformer-based conditional language model, which uses a kind of supervised pre-training to generate protein sequences corresponding to arbitrary combinations of conditioning tags. They don’t mention it in their manuscript, but Elnaggar *et al*. [12] have code for sequence generation from XLNet on their [GitHub repository](https://github.com/agemagician/ProtTrans).

Other deep neural network methods have also been used for protein sequence generation: Costello and Martin [9] as well as Hawkins-Hooker *et al*. [13] created variational autoencoders based on CNN architectures, Riesselman *et al*. [23] used a causal dilated CNN. Another notable generative model for protein sequences was recently published by Russ *et al*. [25]. Russ’ model is not based on a neural network, but is an explicit statistical model of the joint distribution of amino acids at pairs of positions in a multiple sequence alignment of sequences drawn from a single family. Ensembles of sequences produced by Russ’ model have statistical properties nearly identical to the training data while still maintaining diversity. In many ways, Russ’ model represents an apotheosis of pre-deep-learning, family-based statistical protein models, and we use it as a baseline for comparison to our generated sequences.

## 2. Approach

### 2.1. Overview

The major focus of the present work is as a proof-of-concept for a new kind of generative process for protein sequences, Gibbs sampling from BERT models. This generative process is aimed to generate diverse sets of new biologically stable proteins retaining functionally important characteristics of given seed sequences. Selection of generated sequences could potentially be further narrowed down through supervised approaches if experimental data is available for a subset of them, but those methods are beyond the scope of this work. Using Gibbs sampling from BERT models was earlier proposed by Wang and Cho as a means of generating natural language text [27]. The idea behind Gibbs sampling of BERT models is to start with some seed sequence, then iteratively select positions from the sequence, use the masked language prediction task to predict the probability distribution of tokens at those sites, and replace the current tokens with new tokens drawn from the predicted probability distribution, then return the resulting sequence or use it for another round of sampling. Recently, Rao *et al*. [21] used Gibbs sampling from a pre-trained ESM model in a similar way to the method we use in this work, but they used a different evaluation metric, and did not experiment with fine tuning or training from scratch.

As our protein BERT models, we use ESM12, ESM34 [24] and ProtBERT-BFD [12]. We compare these models to sequences generated from a statistical model in a previous work [25], sequences generated by a 12-layer BERT model trained from scratch, sequences generated by 3-gram models, and sequences generated based on a hidden Markov model profile of the target protein family. ESM12, ESM34 were previously pre-trained on a diverse set of 27.1 million sequences [24], and ProtBERT-BFD was previously pretrained on a diverse set 2.1 billion sequences [12]. We further fine-tune ESM12 two different ways: on a set of 994 sequences from the chorismate mutase type II family ([CM_2](http://pfam.xfam.org/family/PF01817.21)), with an additional 226 sequences held out as a test set, and on a set of 2953 sequences from the [Tautomerase](http://pfam.xfam.org/family/PF01361.21) family, with an additional 738 proteins held out as a test set. The CM_2 family was selected because it was the dataset used by Russ *et al*. [25], so by using that dataset we can compare our results to theirs. The CM_2 family is also convenient for testing models because the sequences are relatively short, about 95 amino acids, so training and inference take fewer computational resources than would be needed for longer protein sequences. We selected the Tautomerase family because proteins from that family are also smaller than 100 amino acids, and the family is not at all similar to the CM_2 family, so fine-tuning on one should presumably not improve performance on the other. Our 3-gram models were trained using maximum likelihood objectives on both the CM_2 and the Tautomerase datasets. N-gram/skip-gram models have been demonstrated to effectively generate features for protein classification [3].

We evaluate the quality of the sequences generated by the different models using four different metrics: similarity to the family of the seed sequence as measured by alignment to a hidden Markov model (HMM) of the family [13], similarity to training data as measured by highest alignment score to any sequence from the training set, comparison of first and second order statistics [25, 13] between the generated sequences and the training data, and ability to predict high-performing mutations in the GB-1 protein.

We found that the large pre-trained models, ESM34 and ProtBERT-BFD were able to effectively generate diverse sets sequences that retained functionally important patterns, as measured by scoring against the CM_2 HMM profile. Pre-trained ESM12 produced sequences of much poorer quality than ESM34 and ProtBERT-BFD, but fine-tuning ESM12 allowed generation of sequences whose quality met or surpassed the quality of sequences generated from larger models. The 12-layer BERT model trained from scratch performed similarly to the fine-tuned ESM12. Together these results suggest that Gibbs sampling on protein BERT models could be a powerful tool in helping to guide protein improvement workflows.

### 2.2. Models, Third Party Resources, and Code Availability

All deep learning work was done through the [PyTorch](https://pytorch.org/) framework. Pre-trained ESM models [24] were obtained from a [public repository](https://github.com/facebookresearch/esm). Pre-trained ProtBERT was obtained from the [ProtTrans project](https://github.com/agemagician/ProtTrans) [12]. ProTrans uses the [HuggingFace Transformers library](https://huggingface.co/transformers/), so we took advantage of those interfaces as well. Our 3-gram models were built using the [Natural Language Toolkit for Python](https://www.nltk.org/). Our Gibbs sampler was based on work by Wang and Cho [27], and the associated [code](https://github.com/nyu-dl/bert-gen). HMM generation was done using [hmmemit](http://hmmer.org/) based on an alignment of the same 994 CM_2 sequences used to fine-tune the transformer models. Graphs were produced using Pandas [22], Seaborn [28], and Matplotlib [14]. Our code is available at https://github.com/seanrjohnson/protein_gibbs_sampler.

### 2.3. Data Representations

For each language model variant, sequences had to be processed consistent with how they were during their initial pre-training. For this, each pretrained model provides a tokenizer along with the associated vocabulary that was used for training the original model. For models from the ESM set, we used the tokenizer from ESM source which tokenizes sequences by splitting at the character level, encoding each amino acid independently. ProtBert provides a tokenizer through the transformers package which also tokenizes at the character level. ESM’s tokenizer expects a contiguous string of amino acids for each sequence while Prot-Bert uses a split sequence with a space between each character. During training, batches of tokenized sequences are truncated to a maximum length and any sequence shorter than the max length is padded with a special padding token up to the max length.

### 2.4. Fine Tuning

To evaluate the effect on targeted protein families downstream, additional fine-tuning was performed on ESM12 for each dataset. Fine-tuning was done through self supervised training via the masked language modeling objective where a sample of tokens are corrupted and used for learning. The mechanism for masking is typically a part of pre-training, and was not provided with the ESM models. To further pre-train we wrapped the ESM model and implemented the corruption mechanism following [11] where 15% of tokens are selected at random and masked with 80% probability, swapped with another amino acid randomly with 10% probability, and kept the same with 10% probability. Labels then come from the ground truth amino acids for the positions corrupted during this process while all other amino acids have their label set to a negative integer that is later used by the loss function. The corrupted sequences, along with their labels are then fed through the model which is tasked with predicting the positions selected during the corruption process. After predicting on a batch of sequences, cross entropy loss is calculated between the predictions and target labels for only the corrupted positions, making using of the negative integer in the labels for ignoring all other positions. This loss was used to determine the best model, selecting the model which achieved the lowest loss on the evaluation data prior to over-fitting.

For fine-tuning the ESM12 model, an Adam optimizer was used with parameters (β_1_1=0.9, β_2_=0.999) as used in[24] during initial training of the model. Compared to the original pre-training, we focused on a much smaller dataset. So while the model was originally trained with a learning rate starting from 10e-4, we used a smaller learning rate of 2e-5 to better maintain information learned from the initial pre-training on a much larger dataset. The size of the larger models such as ESM34 also posed a challenge given the hardware we had available to us. While ESM model was trained with 131,072 tokens per batch we only trained with a batch size of 4,096 tokens. Even so, we found our model converged with additional fine-tuning and early stopping.

We also trained a 12-layer BERT model from scratch on the CM_2 data. It has 12 attention heads, a hidden size of 768, and an intermediate size of 3072. It has around 85 million parameters. The model was trained for 100 epochs with a default AdamW optimizer on the MLM objective with a 15% masking probability. We used the pretrained ProtBert tokenizer in conjunction with this model.

### 2.5. Samplers

Drawing from [27], our samplers used a number of modes Table 2: Sequential sampling, where tokens are predicted and added one at a time from left to right across the sequence with new tokens selected randomly according to the predicted probability distribution. Random Sampling, where 10% of positions are selected for sampling each iteration for 20 iterations. Random Sampling with Burn-in, which is the same as Random Sampling except that in the last 10 rounds only the most-likely amino acid is chosen instead of drawing from the distribution. Unless otherwise noted, our starting, ”seed” sequence was the first 95 amino acids of the *E. Coli* [pheA protein](https://www.ncbi.nlm.nih.gov/protein/KRR50589.1), encompassing the CM_2 domain. For some experiments, we used only first 20 amino acids of the CM_2 domain, and for some experiments we used an empty start sequence. When we started with a seed of 20 amino acids, we sampled one position at a time from positions 21 to 95, then wrapped around and started at position one, continuing until each position had been sampled twice. When we started from an empty seed, we did a single pass of sequential sampling. For all sampler experiments detailed here, we generated sequences of length 95.

## 3. Experiments and Results

Evaluating the output quality of a generative model is a nontrivial task. For models that generate human language text, the gold standard is whether a human reader can distinguish whether the text was written by an algorithm or a human. Other, more scalable metrics include statistical measures such as BLEU scores [19] and measuring perplexity of generated sequences as calculated by a model with different architecture. For protein sequences, we consider the gold standard to be whether the protein can be produced and fold into a functional conformation in a biological system. In other words, the best way to judge the quality of a generated protein sequence is to get into the biology lab and do some experiments with it. There are also some statistical measures that can serve as reasonable proxies or baseline measures of protein sequence quality. McGee *et al*. [18] express an alternative perspective. They favor evaluating models by ”generative capacity”, which they define as ”the ability of the model to generate new sequences drawn from the model distribution p(S), which are statistically indistinguishable from those of a given ’target’ protein family.” We contend that trying to match a particular statistical distribution is useful only insofar as the target statistical distribution describes the portion of sequence space that has some desired in-vivo properties. Depending on the target properties, various subsets of natural sequences may only capture a subset of that distribution, or none of it at all in the case of new-to-nature functions. The functional and statistical view of generation quality are not necessarily in opposition, but researchers should be aware of the distinctive perspectives.

Previous work on protein generative models have used a variety of different methods to evaluate the quality of generated sequences Table 1. Costello *et al*. [9] simply measured the ability of their autoencoder to reproduce the input sequence. Madani *et al*. [17] use accuracy of predicted secondary structure, stability predicted from physical simulations, similarity to a reference sequence, and ability to select beneficial mutations for a protein called GB-1 for which exhaustive experimental data is available. Riesselman *et al*. [23] use basic statistics such as length and averages over amino acid properties, as well as sequence similarity to training data, they also briefly mention that they did preliminary lab experiments, but do not present any data. Russ *et al*. [25] compare first order sequence statistics and second and third order correlations between their training set and generated proteins, they also calculate sequence similarity between their generated sequences and their training sequences and perform extensive validation experiments in the laboratory. Hawkins-Hooker *et al*. [13] calculated HMM-profile scores, compare first- and second-order sequence statistics, co-variance (a function of second-order statistics), and direct-coupling analysis (a different function of second-order statistics) between training sequences and generated sequences, and perform laboratory experiments. McGee *et al*. [18] used Hamming distance to the training set, second order correlations, approximations of third and higher order correlations, and statistical energy. Rao *et al*. [21] used a multiple sequence alignment of generated sequences to predict contact maps, using direct coupling analysis.

**Table 1.** Protein generation evaluation metrics from previous works.

| Evaluation metric | Costello et al. (2019) | Madani et al. (2020) | Russ et al. (2020) | Riesselman et al. (2019) | Hawkins-Hooker et al. (2020) | McGee et al. (2020) | Rao et al. (2020) | This work |
| --- | --- | --- | --- | --- | --- | --- | --- | --- |
| Sequence similarity to input | X |  |  |  |  |  |  |  |
| Length and amino acid properties |  |  |  | X |  |  |  |  |
| Sequence similarity to training set |  | X | X | X |  | X |  | X |
| HMM profile score |  |  |  |  | X |  |  | X |
| First order sequence statistics |  |  | X |  | X |  |  | X |
| Second order sequence statistics |  |  |  |  | X |  |  | X |
| Covariance (second order correlations) |  |  | X |  | X | X |  | X |
| Third or higher order correlations |  |  | X |  |  | X |  |  |
| direct-coupling analysis |  |  |  |  | X |  | X |  |
| Perplexity/statistical energy |  | X |  |  |  | X |  |  |
| Secondary structure prediction |  | X |  |  |  |  |  |  |
| Stability simulation |  | X |  |  |  |  |  |  |
| GB-1 prediction |  | X |  |  |  |  |  | X |
| Experimental characterization |  |  | X | ? |  |  |  |  |

**Table 2.**
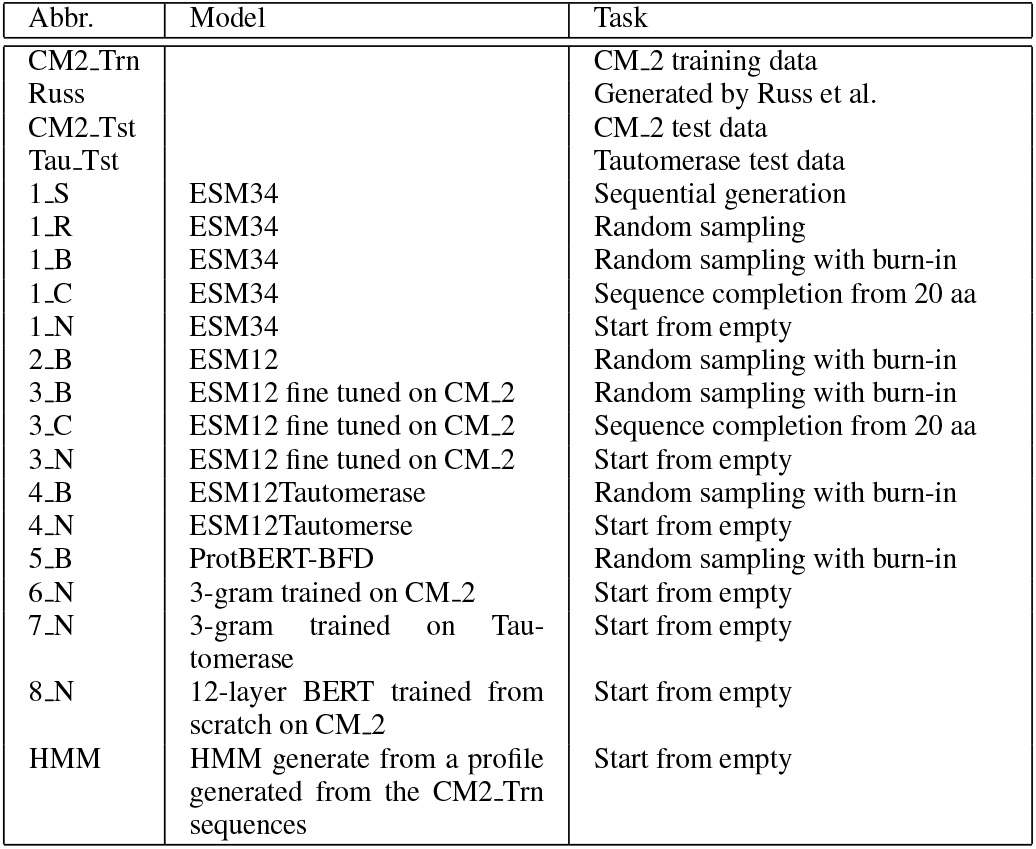
Summary of models and sampling conditions

In the present work, we measure similarity to the training set, HMM profile scores, first- and second-order statistics, second-order correlations, and prediction of GB-1 variants. We follow Russ *et al*. [25] in focusing on the (CM_2) family so that we can compare to their results.

### 3.1. HMM profile score

HMM profile score (Figure 1, Top) is a family-specific statistical profile indicating whether a sequence contains amino acids that are critical for function of a protein in a particular family, calculated using the [HMMER package](http://hmmer.org/). A protein may have low overall similarity to other proteins in a family but keep important amino acid arrangements and still have the same function. ESM34 and ProtBert do a remarkable job of generating sequences that have high HMM scores for CM_2 when provided a full-length seed, but fail when seeded with 20 amino acids or shorter. Sequences generated from pre-trained ESM12 when given a full-length seed score somewhat worse than ESM34. ESM12 finetuned on CM_2 sequences generates sequences matching the CM_2 profile even when given no seed at all. ESM12 fine-tuned on Tautomerases produces sequences matching the Tautomerase HMM even when seeded with a CM_2 protein. The 12-layer BERT model trained from scratch generates sequences that match the CM_2 profile well. 3-gram model controls perform poorly.

**Figure 1.**
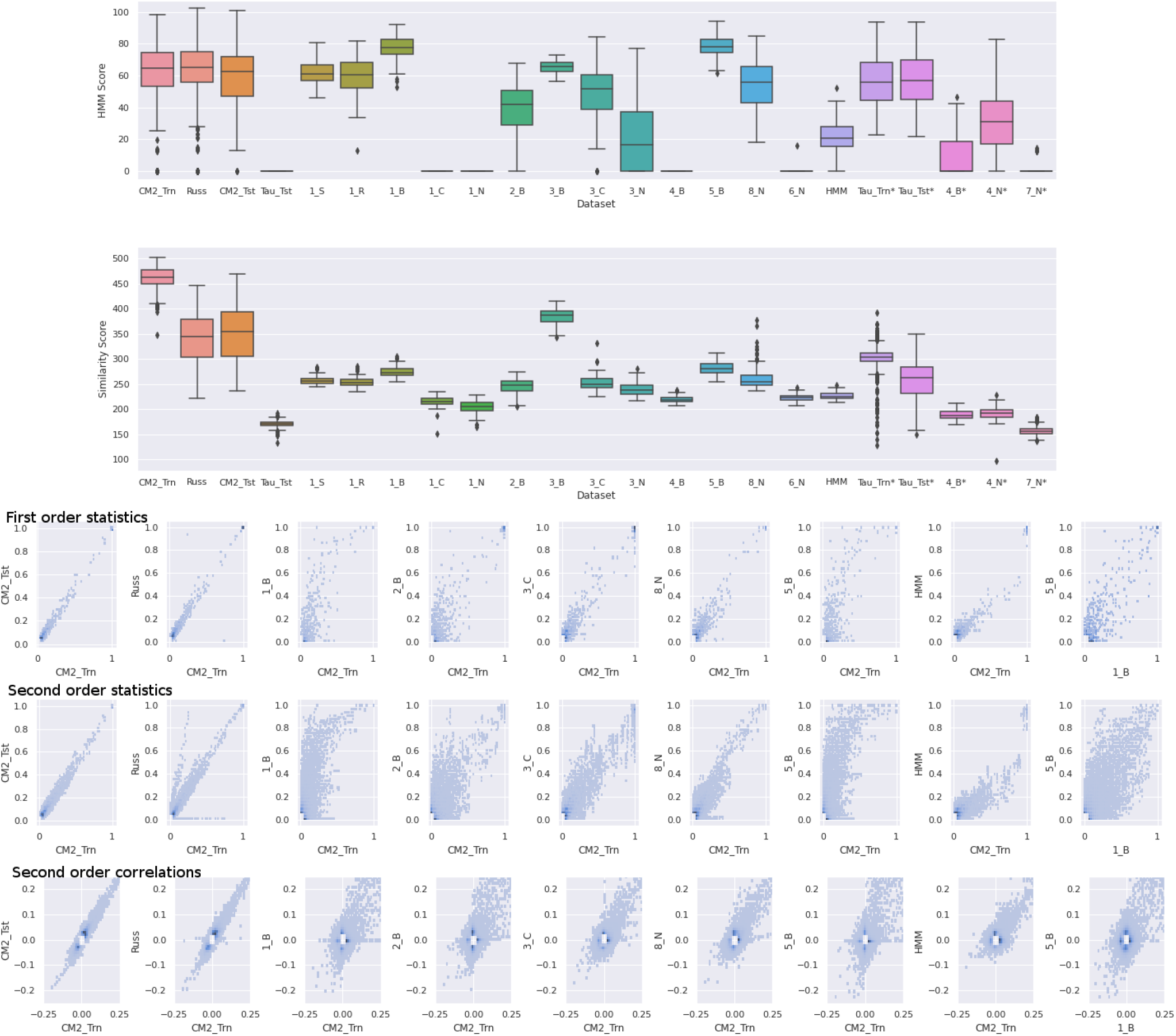
Top: HMM profile scores. Sequences are scored against CM_2 family profile except for conditions marked with a *, which are scored against the Tautomerase family profile. For reference datasets, all sequences in the dataset were used. For generated datasets, 100 sequences were generated per condition. Middle: Alignment scores to best match from the training set. Datasets marked with an * were compared to the Tautomerase training dataset, all others were compared to the CM_2 training dataset. For reference datasets, all sequences in the dataset were used. For generated datasets, 100 sequences were generated per condition. Bottom: Comparison of first order statistics (top), second order statistics (middle), and second order correlations (bottom) between multiple sequence alignments of various datasets. Points near the origin omitted for clarity.

### 3.2. Alignment to training set

To check whether the high HMM profile scores for finetuned models were due to memorizing training sequences, we aligned the generated sequences with the training sequences. The expectation is that the generated sequences would have some similarities, but not be identical to the training sequences. We used the alignment calculation functionality from the nwalign3 library to find the training sequence that had the highest alignment score with each generated sequence. We used the BLOSUM62 scoring matrix with a gap open penalty of 0.5 and a gap extension penalty of 0.1 for internal gaps, and 0 and 0 for end gaps. As a benchmark, we also performed the analysis on the training and test datasets. We found (Figure 1, Middle) that with the exception of fine-tuned ESM12 with a full-length seed (3_B), the transformer models produced a wide diversity of proteins, and are clearly not just memorizing training data, despite producing good HMM profile scores and second order statistics. It should be noted that some of the models were not fine-tuned (such as ESM34 and ProtBert-BFD), so it would be better to compare sequences generated by those models to the larger corpus of the UniProt or BFD databases which they were pre-trained on. We did not systematically evaluate those similarities, but we did spot check a few generated sequences by BLAST searches against the UniProt database and saw top hits of around of 50% identity, which is consistent with our conclusion that over-fitting is not a major concern for sequence generation from these models; the models appear to produce a wide diversity of sequences that are not present in the training data, but still have good HMM scores.

### 3.3. First and second order statistics

First order statistics are a comparison of the frequency of each amino acid at each position between two datasets, for example a point at (0.5,0.5) would mean that for some position there is an amino acid which occurs in 50% of the generated sequences and 50% of the reference sequences. Second order statistics are similar but involve two positions instead of one: a point at (0.5,0.3) could indicate that 30% of the generated sequences have *both* a ”G” at position 5 *and* an ”A” at position 7, while that combination occurs in 50% of reference sequences. Interactions between distant amino acids can be critical for protein stability and function, so sets of generated proteins with second order statistics similar to natural sequences are likely to be stable and functional [25, 13]. To calculate these statistics, we first used [MAFFT](https://mafft.cbrc.jp/alignment/software/), using options ’–globalpair–maxiterate 1000’ to create multiple sequence alignments of all sequences for each pair of datasets (the x-axis dataset and y-axis dataset), then tabulated the frequencies within each dataset. Second order correlations are a function of first and second order statistics: for a pair of amino acids and positions, the correlation is equal to the second order statistic minus the product of the first order statistics [25].

We see (Figure 1, Bottom) that the CM_2 test set closely matches the statistics of the training data, and that many of our models, including pre-trained ESM34 (1_B) and ESM12 (2_B), and fine-tuned ESM12 (3_C) are able to approximate the first order statistics of the reference data. Fine-tuned ESM12 (3_C) does almost as well as the sequences form Russ *et al*. [25] at matching even the second order statistics of the training data, even though it was not specifically trained to do that and we were not optimizing hyper-parameters with that in mind. We also observed that a 12-layer BERT model trained from scratch (8_N), given no seed, performs similarly to fine-tuned ESM12 model given a 20 sequence seed.

It is notable that the graphs for ESM34 and ProtBert look remarkably similar, even though ESM34 has more parameters and ProtBERT was trained on more than 10 times as many sequences. It is also notable that those models don’t match the reference data very well, particularly in terms of second order statistics. This is not too surprising, considering that they were not specifically fine-tuned on CM_2_Trn, but it does raise the question of whether the part of sequence space that they are generating in is as rich with stable, functional proteins as the space occupied by the reference sequences. It could be that training on huge protein databases give ESM34 and ProtBert access to an equally stable area of protein space that is not reachable by models trained only on single family or models heavily fine-tuned on a single family. It would be very interesting to test this hypothesis with laboratory experiments.

### 3.4. GB-1 protein generation

We replicated an experiment performed by Madani *et al*. [17] to validate the fitness of the models by predicting GB-1 variants. The models did not perform better than randomly generated variants, see appendix for details.

### 3.5. Future Directions

We did not put a great deal of effort into finding optimal hyperparmeters for fine-tuning or sampling, so the simplest way to improve the quality of generated sequences may just be to do hyperparameter scans. Deciding on an objective function for the scans could be difficult, but a good place to start might be to try to maximize second order correlations between generated sequences and training sequences.

Compared to Direct Coupling Analysis-based approaches [25], ESM34 and ProtBERT may be able to predict stable mutations in de-novo proteins or those where not much relevant information is known. We could not test this, because we don’t have access to the complete training data for ESM, so we don’t know what the held-out protein families were. Large pre-trained transformer models or RNNs may also have access to stable parts of sequence space that cannot be inferred from single family multiple sequence alignments; recent experimental results from Biswas *et al*. [5] support this idea.

Another interesting future direction could be to examine the generative capacity of small transformer models, even smaller than ESM12. Results from a recent paper [4] give hope that this might be possible. Fine-tuning only the last layer of ESM34 or ProtBERT might also improve performance on a target family without needing significant compute power for the back propagation. Alternative techniques for generating sequences may also be explored. For example, a fine-tuned BERT model may be leveraged to teach a smaller auto-regressive model via knowledge distillation. This approach has been shown to work well on text generation tasks [8]. Going the other way, recently, the GPT-3 model, with 175 billion trainable parameters, has shown impressive natural language text generation capabilities [7], raising the possibility that simply scaling existing protein LMs might also lead to better generative performance.

A major limitation of both ProtBERT and ESM for sequence generation tasks is that neither model was pretrained with masking of the end-of-sequence token, so the Gibbs sampler doesn’t know when to stop predicting new amino acids. This means that the sequence length must be provided as a hyperparameter, and all generated sequences will be of that fixed length. Pre-training could be made more amenable to Gibbs sampling by including an end-of sequence token in the MLM pre-training. Another possibility would be to incorporate token-deletion and text-infilling tasks, such as those used in training of the BART model [16], to allow the sampler to dynamically expand or contract the sequence. A simple variation of token-deletion and text-infilling would be to include a zero-length amino acid token in the vocabulary and randomly insert the new token into sequences before pre-training.

## 4. Acknowledgments

We acknowledge the help and support of Siddharth Goyal. This work was done as a class project for CS 7643: Deep Learning as part of the Online Master of Science in Computer Science (OMS CS) program at the Georgia Institute of Technology.

## 5. Appendix

### 5.1. GB-1 protein generation

To validate the model is generating stable proteins, an experiment first performed by Madani *et al*. [17] has been modified and replicated. The experiment analyzes how well the model generates specific amino acids in protein G domain B1 (GB1). The amino acids in four specific positions have been previously studied and categorized with a fitness score that reflects their stability and function. The dataset contains 149,361 of the possible 160,000 combinations of these four amino acids as well as their scores. In the evaluation performed by Madani *et al*., they generated perplexity scores for every known combination, and then looked for a correlation between the perplexity and the fitness score. We focused our evaluation on the ability of the sampler to generate those four values. The four positions were masked, then the sampler re-generated these values in parallel. The models that were compared were ESM34, ESM12, and the two fine-tuned ESM models.

The results of 300 iterations for each model are shown in Table 3. Although the max values are very high, these are outliers and could easily have been selected at random. It appears that the ESM12 model improved when fine-tuned with the CM_2 data. All other models do not seem to outperform random selection. Since the two families are not related, this result seems to be an artifact. Our result differs from the that of Madani *et al*. [17], who saw an improvement over the random selection. This could be due to the fact that they had selected the top 100 sequences by perplexity, whereas we used all generated samples.

**Table 3.** Summarized results of the GB1 experiment

| Model | Min | Max | Mean |
| --- | --- | --- | --- |
| ESM34 | 0.0 | 3.35696 | 0.08195 |
| ESM12 | 0.0 | 5.39813 | 0.09020 |
| ESM12 - CM2 | 0.0 | 3.93922 | 0.13656 |
| ESM12 - Tautomerase | 0.0 | 4.03464 | 0.08099 |
| Random | 0.0 | 3.22443 | 0.11015 |

## Notes

### Competing Interest Statement

SJ is currently employed by Conagen Inc., a synthetic biology company. This work was not sponsored or overseen by Conagen Inc. in any way.

## References

[1] Ethan C. Alley, Grigory Khimulya, Surojit Biswas, Mohammed AlQuraishi, and George M. Church. Unified rational protein engineering with sequence-based deep representation learning. Nature Methods, 16(12):1315–1322, Dec. 2019. Number: 12 Publisher: Nature Publishing Group. 2

[2] Frances H. Arnold. Directed evolution: Bringing new chemistry to life. Angewandte Chemie International Edition, 57(16):4143–4148, 2018. _eprint: https://onlinelibrary.wiley.com/doi/pdf/10.1002/anie.201708408. 1

[3] Christopher Michel Kearney Ashiqul Islam, Benjamin J Heil and Erich J Bakerb. Protein classification using modified ngram and skip-gram models. bioRxiv, 2017. 3

[4] Nicholas Bhattacharya, Neil Thomas, Roshan Rao, Justas Dauparas, Peter K. Koo, David Baker, Yun S. Song, and Sergey Ovchinnikov. Single Layers of Attention Suffice to Predict Protein Contacts. bioRxiv, page 2020.12.21.423882, Dec. 2020. Publisher: Cold Spring Harbor Laboratory Section: New Results. 8

[5] Surojit Biswas, Grigory Khimulya, Ethan C. Alley, Kevin M. Esvelt, and George M. Church. Low-N protein engineering with data-efficient deep learning. bioRxiv, page 2020.01.23.917682, Aug. 2020. Publisher: Cold Spring Harbor Laboratory Section: New Results. 2, 8

[6] Uwe T. Bornscheuer, Bernhard Hauer, Karl Erich Jaeger, and Ulrich Schwaneberg. Directed evolution empowered redesign of natural proteins for the sustainable production of chemicals and pharmaceuticals. Angewandte Chemie International Edition, 58(1):36–40, 2019. _eprint: https://onlinelibrary.wiley.com/doi/pdf/10.1002/anie.201812717. 1

[7] Tom B. Brown, Benjamin Mann, Nick Ryder, Melanie Subbiah, Jared Kaplan, Prafulla Dhariwal, Arvind Neelakantan, Pranav Shyam, Girish Sastry, Amanda Askell, Sandhini Agarwal, Ariel Herbert-Voss, Gretchen Krueger, Tom Henighan, Rewon Child, Aditya Ramesh, Daniel M. Ziegler, Jeffrey Wu, Clemens Winter, Christopher Hesse, Mark Chen, Eric Sigler, Mateusz Litwin, Scott Gray, Benjamin Chess, Jack Clark, Christopher Berner, Sam McCandlish, Alec Radford, Ilya Sutskever, and Dario Amodei. Language Models are Few-Shot Learners. arXiv:2005.14165 [cs], July 2020. arXiv: 2005.14165. 8

[8] Yen-Chun Chen, Zhe Gan, Yu Cheng, Jingzhou Liu, and Jingjing Liu. Distilling knowledge learned in bert for text generation. arXiv, 2020. 8

[9] Zak Costello and Hector Garcia Martin. How to hallucinate functional proteins. arXiv:1903.00458 [q-bio], 2019. 3, 5

[10] Zihang Dai, Zhilin Yang, Yiming Yang, Jaime Carbonell, Quoc V. Le, and Ruslan Salakhutdinov. Transformer-XL: Attentive language models beyond a fixed-length context. arXiv:1901.02860 [cs, stat], 2020. 2

[11] Jacob Devlin, Ming-Wei Chang, Kenton Lee, and Kristina Toutanova. BERT: Pre-training of deep bidirectional transformers for language understanding. arXiv:1810.04805 [cs], 2020. 2, 4

[12] Ahmed Elnaggar, Michael Heinzinger, Christian Dallago, Ghalia Rehawi, Yu Wang, Llion Jones, Tom Gibbs, Tamas Feher, Christoph Angerer, Martin Steinegger, Debsindhu Bhowmik, and Burkhard Rost. ProtTrans: Towards cracking the language of life’s code through self-supervised deep learning and high performance computing. bioRxiv, page 2020.07.12.199554, 2020. Publisher: Cold Spring Harbor Laboratory Section: New Results. 2, 3

[13] Alex Hawkins-Hooker, Florence Depardieu, Sebastien Baur, Guillaume Couairon, Arthur Chen, and David Bikard. Generating functional protein variants with variational autoencoders. bioRxiv, 2020. 3, 5, 6

[14] J. D. Hunter. Matplotlib: A 2D Graphics Environment. Computing in Science Engineering, 9(3):90–95, May 2007. Conference Name: Computing in Science Engineering. 4

[15] Zhenzhong Lan, Mingda Chen, Sebastian Goodman, Kevin Gimpel, Piyush Sharma, and Radu Soricut. ALBERT: A lite BERT for self-supervised learning of language representations. arXiv:1909.11942 [cs], 2020. 2

[16] Mike Lewis, Yinhan Liu, Naman Goyal, Marjan Ghazvininejad, Abdelrahman Mohamed, Omer Levy, Veselin Stoyanov, and Luke Zettlemoyer. BART: Denoising Sequence-to-Sequence Pre-training for Natural Language Generation, Translation, and Comprehension. In Proceedings of the 58th Annual Meeting of the Association for Computational Linguistics, pages 7871–7880, Online, 2020. Association for Computational Linguistics. 8

[17] Ali Madani, Bryan McCann, Nikhil Naik, Nitish Shirish Keskar, Namrata Anand, Raphael R. Eguchi, Po-Ssu Huang, and Richard Socher. ProGen: Language modeling for protein generation. arXiv:2004.03497 [cs, q-bio, stat], 2020. 2, 5, 7, 10

[18] Francisco McGee, Quentin Novinger, Ronald M. Levy, Vincenzo Carnevale, and Allan Haldane. Generative Capacity of Probabilistic Protein Sequence Models. Dec. 2020. 5

[19] Kishore Papineni, Salim Roukos, Todd Ward, and Wei-Jing Zhu. Bleu: a Method for Automatic Evaluation of Machine Translation. In Proceedings of the 40th Annual Meeting of the Association for Computational Linguistics, pages 311–318, Philadelphia, Pennsylvania, USA, July 2002. Association for Computational Linguistics. 4

[20] Roshan Rao, Nicholas Bhattacharya, Neil Thomas, Yan Duan, Peter Chen, John Canny, Pieter Abbeel, and Yun Song. Evaluating Protein Transfer Learning with TAPE. Advances in Neural Information Processing Systems, 32:9689–9701, 2019. 2

[21] Roshan M. Rao, Joshua Meier, Tom Sercu, Sergey Ovchinnikov, and Alexander Rives. Transformer protein language models are unsupervised structure learners. bioRxiv, page 2020.12.15.422761, Dec. 2020. Publisher: Cold Spring Harbor Laboratory Section: New Results. 3, 5

[22] Jeff Reback, Wes McKinney, jbrockmendel, Joris Van den Bossche, Tom Augspurger, Phillip Cloud, gfyoung, Sinhrks, Adam Klein, Matthew Roeschke, Simon Hawkins, Jeff Tratner, Chang She, William Ayd, Terji Petersen, Marc Garcia, Jeremy Schendel, Andy Hayden, MomIsBestFriend, Vytautas Jancauskas, Pietro Battiston, Skipper Seabold, chris b1, h vetinari, Stephan Hoyer, Wouter Overmeire, alimcmaster1, Kaiqi Dong, Christopher Whelan, and Mortada Mehyar. pandas-dev/pandas: Pandas 1.0.3, Mar. 2020. 4

[23] Adam Riesselman, Jung-Eun Shin, Aaron Kollasch, Conor McMahon, Elana Simon, Chris Sander, Aashish Manglik, Andrew Kruse, and Debora Marks. Accelerating protein design using autoregressive generative models. bioRxiv, page 757252, 2019. Publisher: Cold Spring Harbor Laboratory Section: New Results. 3, 5

[24] Alexander Rives, Joshua Meier, Tom Sercu, Siddharth Goyal, Zeming Lin, Demi Guo, Myle Ott, C. Lawrence Zitnick, Jerry Ma, and Rob Fergus. Biological structure and function emerge from scaling unsupervised learning to 250 million protein sequences. bioRxiv, page 622803, 2020. Publisher: Cold Spring Harbor Laboratory Section: New Results. 2, 3, 4

[25] William P. Russ, Matteo Figliuzzi, Christian Stocker, Pierre Barrat-Charlaix, Michael Socolich, Peter Kast, Donald Hilvert, Remi Monasson, Simona Cocco, Martin Weigt, and Rama Ranganathan. An evolution-based model for designing chorismate mutase enzymes. Science, 369(6502):440–445, 2020. Publisher: American Association for the Advancement of Science Section: Report. 3, 5, 6, 8

[26] Martin Steinegger, Milot Mirdita, and Johannes Soding. Protein-level assembly increases protein sequence recovery from metagenomic samples manyfold. Nature Methods, 16(7):603–606, July 2019. Number: 7 Publisher: Nature Publishing Group. 1

[27] Alex Wang and Kyunghyun Cho. BERT has a mouth, and it must speak: BERT as a markov random field language model. arXiv:1902.04094 [cs], 2020. 3, 4

[28] Michael Waskom, Olga Botvinnik, Maoz Gelbart, Joel Ostblom, Paul Hobson, Saulius Lukauskas, David C Gemperline, Tom Augspurger, Yaroslav Halchenko, Jordi Warmenhoven, John B. Cole, Julian de Ruiter, Jake Vanderplas, Stephan Hoyer, Cameron Pye, Alistair Miles, Corban Swain, Kyle Meyer, Marcel Martin, Pete Bachant, Eric Quintero, Gero Kunter, Santi Villalba, Brian, Clark Fitzgerald, C.G. Evans, Mike Lee Williams, Drew O’Kane, Tal Yarkoni, and Thomas Brunner. mwaskom/seaborn: v0.11.0 (Sepetmber 2020), Sept. 2020. 4

[29] Jianyi Yang, Ivan Anishchenko, Hahnbeom Park, Zhenling Peng, Sergey Ovchinnikov, and David Baker. Improved protein structure prediction using predicted interresidue orientations. Proceedings of the National Academy of Sciences, 117(3):1496–1503, 2020. Publisher: National Academy of Sciences Section: Biological Sciences. 2

[30] Zhilin Yang, Zihang Dai, Yiming Yang, Jaime Carbonell, Ruslan Salakhutdinov, and Quoc V. Le. XLNet: Generalized Autoregressive Pretraining for Language Understanding. arXiv:1906.08237 [cs], Jan. 2020. arXiv: 1906.08237. 2

